# Positionally distinct interferon stimulated dermal immune acting fibroblasts promote neutrophil recruitment in Sweet’s syndrome

**DOI:** 10.1101/2024.06.24.600500

**Authors:** Kellen J. Cavagnero, Julie Albright, Fengwu Li, Tatsuya Dokoshi, Rachael Bogle, Joseph Kirma, J. Michelle Kahlenberg, Allison C. Billi, Jennifer Fox, Anthony Coon, Craig J. Dobry, Brian Hinds, Lam C. Tsoi, Paul W. Harms, Johann E. Gudjonsson, Richard L. Gallo

## Abstract

Sweet’s syndrome is a poorly understood inflammatory skin disease characterized by neutrophil infiltration to the dermis. Single-nucleus and bulk transcriptomics of archival clinical samples of Sweet’s syndrome revealed a prominent interferon signature in Sweet’s syndrome skin that was reduced in tissue from other neutrophilic dermatoses. This signature was observed in different subsets of cells, including fibroblasts that expressed interferon-induced genes. Functionally, this response was supported by analysis of cultured primary human dermal fibroblasts that were observed to highly express neutrophil chemokines in response to activation by type I interferon. Furthermore, single-molecule resolution spatial transcriptomics of skin in Sweet’s syndrome identified positionally distinct immune acting fibroblasts that included a CXCL1+ subset proximal to neutrophils and a CXCL12+ subset distal to the neutrophilic infiltrate. This study defines the cellular landscape of neutrophilic dermatoses and suggests dermal immune acting fibroblasts play a role in the pathogenesis of Sweet’s syndrome through recognition of type I interferons.

## Introduction

Sweet’s syndrome (SS) is an uncommon inflammatory skin disease characterized by painful red nodules or plaques with dense neutrophil infiltrate on the face, neck, or arms (Joshi et al., 2022). Therefore, SS belongs to the class of neutrophilic dermatoses, which also includes pyoderma gangrenosum (PG), pustular psoriasis (PP), and Behcet disease. Although SS is often idiopathic, it can be drug-induced and has been associated with various immune-related conditions such as cancer, infection, inflammatory diseases, vaccination, and pregnancy. The etiology of SS remains largely unknown but is thought to involve aberrant neutrophils (Bhattacharya et al., 2023), genetic factors (Bhattacharya et al., 2023), and proinflammatory cytokines such as IL-1, TNFα, and IL-6 (Joshi et al., 2022, Heath and Ortega-Loayza, 2019, Bhattacharya et al., 2023). While systemic steroids effectively treat many SS patients, there is a pressing need for novel therapeutic approaches to address steroid resistance and minimize side effects.

Recent technological advancements have enabled detailed transcriptomic analysis of fresh and formalin-fixed paraffin-embedded (FFPE) tissue samples, providing an opportunity to comprehensively characterize gene expression during disease (Guo et al., 2023). Such analysis has advanced understanding of the pathophysiology of many diseases ranging from atopic dermatitis (He et al., 2020) to inflammatory bowel disease (Smillie et al., 2019) to Alzheimer’s disease (Mathys et al., 2019). Due to the difficulty of obtaining fresh SS tissue, a detailed characterization of gene expression in SS has not been reported.

Here, we define the skin cellular and molecular landscape of a large cohort of patients with SS, PG, and PP using cutting-edge single-nucleus RNA sequencing (snRNA-Seq), bulk RNA sequencing (bulk RNA-Seq), and subcellular resolution spatial transcriptomics approaches. Integrated analysis identified a prominent and unique type I interferon (IFN) signature in SS in various subsets of cells including fibroblasts. Functional experiments with primary human dermal fibroblasts demonstrated that type I IFN activated these cells to highly express inflammatory mediators relevant to SS including neutrophil chemokines. Overall, this study reveals a hitherto unknown role for type I IFN, through activation of immune acting fibroblasts, to drive neutrophil inflammation in SS.

## Results

### Activation by interferon is a prominent feature of several Sweet’s syndrome skin cell subsets including fibroblasts

To understand the cellular landscape of SS and other neutrophilic dermatoses, we employed single-nucleus transcriptomics using the 10X Genomics Flex platform of archived clinical formalin-fixed paraffin-embedded (FFPE) lesional skin biopsies from patients with SS (n=6). A similar analysis was also performed on other neutrophilic disorders, (pyoderma gangrenosum (PG, early-stage disease, n=6), and pustular psoriasis (PP, n=1), as well as healthy controls (HC, n=5), to identify potentially unique gene signatures within SS using the 10X Genomics Flex platform (Figure 1A). In total, we recovered 74,563 nuclei, with an average of 1036 genes and 1621 unique transcripts detected per nucleus (Figure S1A). Quality control pseudo-bulk principal component analysis confirmed that neutrophilic dermatosis samples were transcriptionally distinct from HC (Figure S1B).

**Figure 1:**
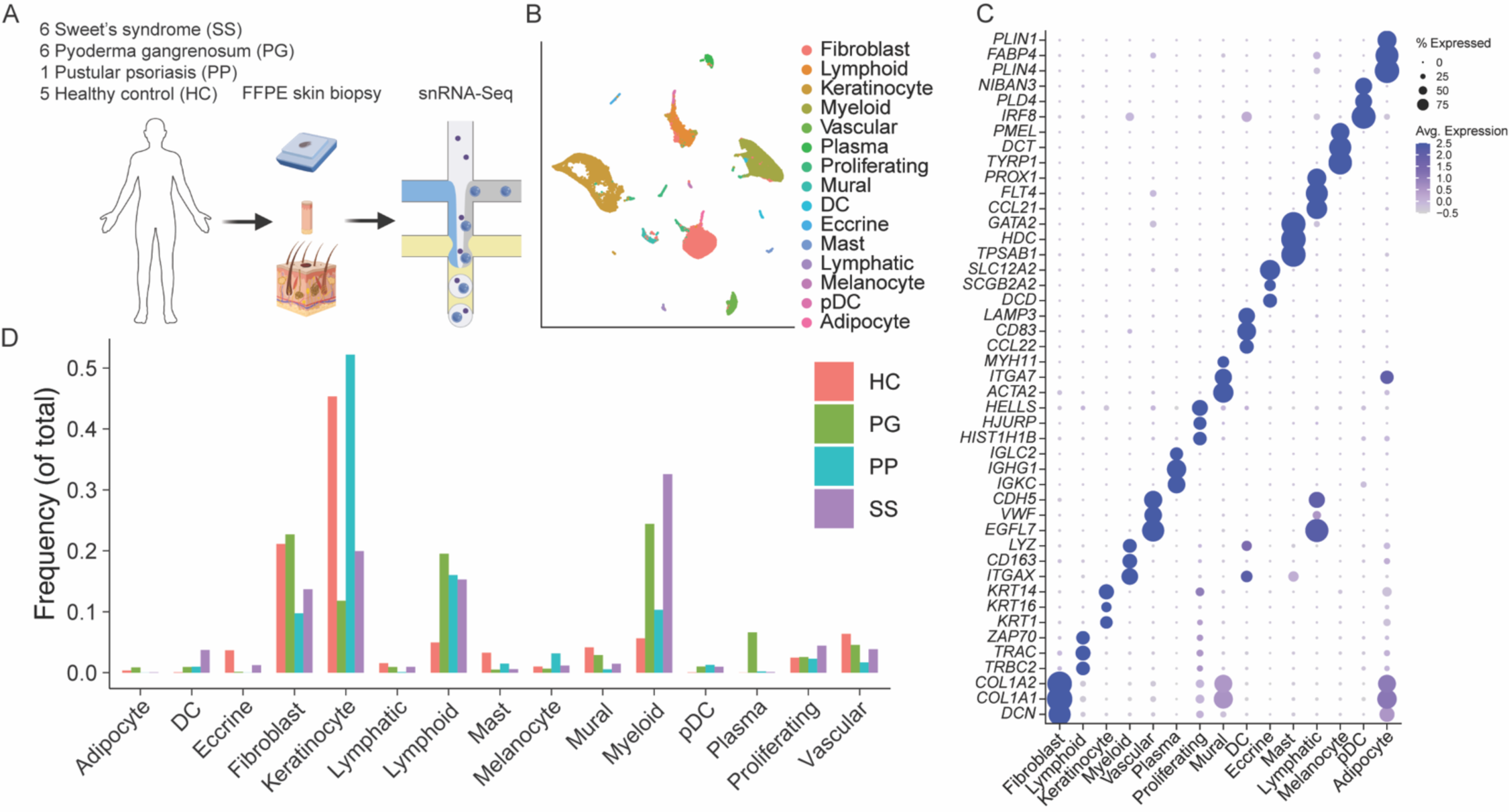
Single-nucleus RNA sequencing of neutrophilic dermatoses. (A) Experimental schematic. (B) Dimensionality reduction and unsupervised clustering, colored by cell type. (C) Expression of top 3 marker genes per cell type. (D) Proportion of each cell type across conditions. SS, Sweet’s syndrome; PG, pyoderma gangrenosum; PP, pustular psoriasis; HC, healthy control.

Recovered nuclei were annotated using known marker genes following dimensionality reduction and unsupervised clustering (Figure 1B), revealing 15 major cell types (Figure 1C): fibroblasts (*COL1A1, COL1A2, DCN)*, lymphoid cells (*TRAC, TRBC2, ZAP70)*, keratinocytes (*KRT14, KRT16, KRT1)*, myeloid cells (*ITGAX, CD163, LYZ)*, vascular endothelial cells (*EGFL7, VWF, CDH5)*, plasma cells (*IGKC, IGHG1, IGLC2)*, proliferating cells (*HIST1H1B, HJURP, HELLS)*, mural cells (*ITGA7, MYH11, ACTA2)*, dendritic cells (DCs) (*CCL22, CD83, LAMP3)*, eccrine cells (*DCD, SCGB2A2, SLC12A2)*, mast cells (*TPSAB1, HDC, GATA2)*, lymphatic endothelial cells (*CCL21, FLT4, PROX1)*, melanocytes (*DCT, TYRP1, PMEL)*, plasmacytoid dendritic cells (pDCs) (*NIBAN3, PLD4, IRF8*), and adipocytes (*PLIN1, FABP4, PLIN4*). Proliferating cells, lymphoid cells, DCs, pDCs, and myeloid cells—a lineage that includes neutrophils— were enriched in neutrophilic dermatoses lesions compared to HC skin (Figure 1D).

To understand which cell types may drive skin inflammation in SS, we determined the extent of transcriptional response of each cell type by performing differential expression analysis between HC and SS skin (Figure 2A). This analysis revealed that the most transcriptionally activated cell type in SS was fibroblasts, followed by myeloid cells then keratinocytes.

**Figure 2:**
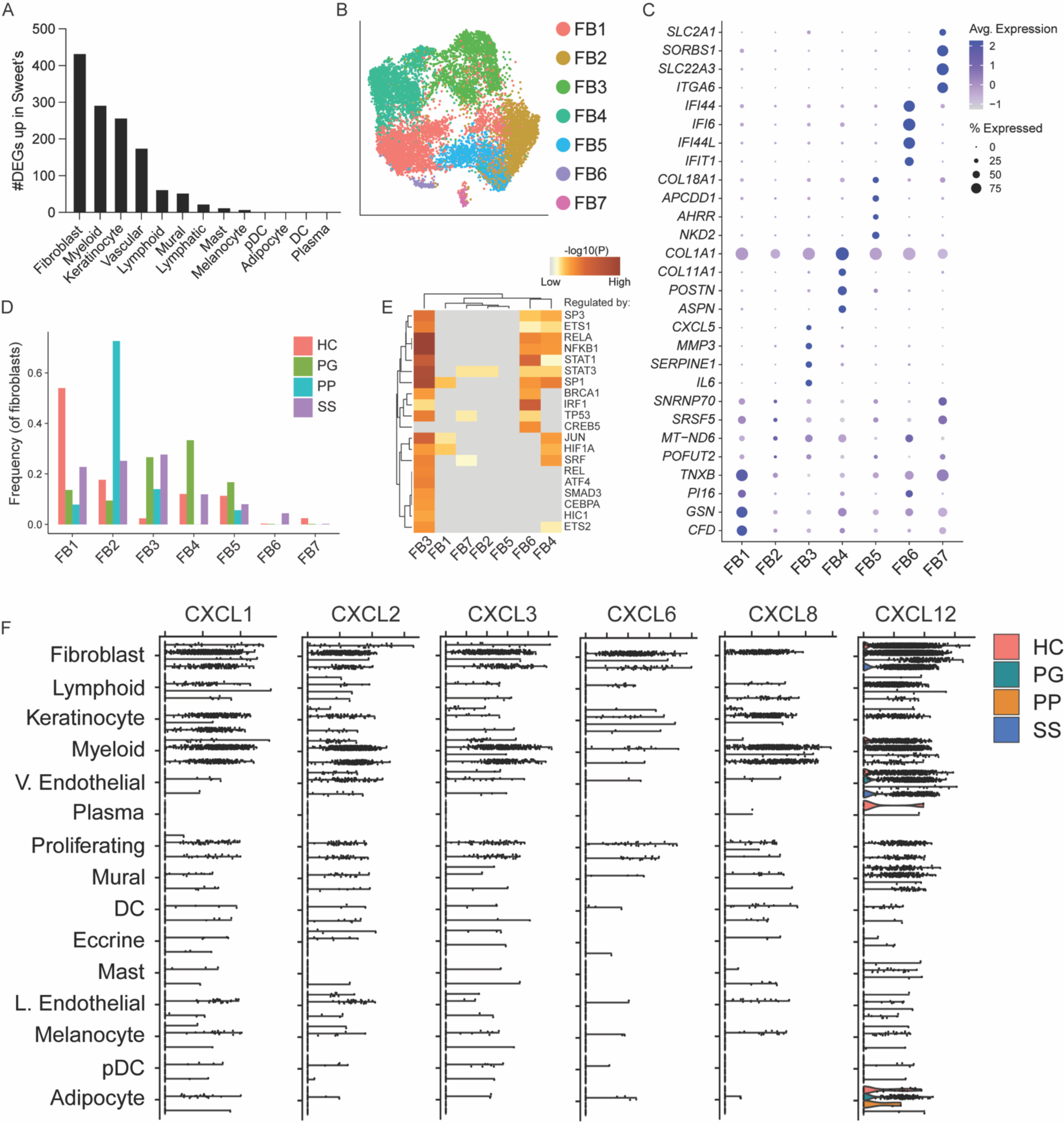
IFN-activated fibroblasts are enriched in Sweet’s syndrome. (A) Number of significantly upregulated genes (P_adj_ < 0.05) in SS compared to HC for each cell type. Differential expression analysis was performed using DESeq2 with pseudobulked data to control for patient heterogeneity. (B) Dimensionality reduction and unsupervised clustering of fibroblasts (FB), colored by subset. (C) Expression of top 4 marker genes per fibroblast subset. (D) Proportion of fibroblast subsets across conditions. (E) Pathway analysis inferring upstream transcription factors regulating gene expression for each cluster. (F) Expression of neutrophil chemokine CXCR2 ligands (*CXCL1, CXCL2, CXCL3, CXCL5, CXCL6, CXCL8*) and CXCR4 ligand (*CXCL12*) across cell types for each condition. SS, Sweet’s syndrome; PG, pyoderma gangrenosum; PP, pustular psoriasis; HC, healthy control; IFN, interferon.

Dimensionality reduction and unsupervised clustering of fibroblasts identified 8 transcriptionally distinct fibroblast subsets (Figure 2B). FB3 was enriched in neutrophilic dermatoses relative to HC and was marked by expression of neutrophil chemokines and proinflammatory cytokines (*CXCL5*, *IL6*) (Figure 2D). FB4 highly expressed markers of extracellular matrix production and myofibroblasts (*COL1A1, COL1A11*) and was uniquely expanded in PG. FB6, marked by interferon (IFN)-induced genes (*IFI44, IFI6, IFI44L, IFIT1*), was found almost exclusively in SS. FB1 highly expressed the universal fibroblast gene *PI16* (Buechler et al., 2021) and was decreased across disease conditions, hinting that this subset may serve as a progenitor for neutrophilic dermatoses-associated fibroblast subsets. Pathway analysis suggested that FB3 and FB6 were regulated by signaling intermediates IFN response factor 1 (IRF1) and STAT1; therefore, these subsets may both be induced by IFN (Figure 2E).

Compared to other cell lineages, fibroblasts in SS and PG were a dominant source of neutrophil chemokine ligands that bind CXCR2 (*CXCL1, CXCL2, CXCL3, CXCL5, CXCL6, CXCL8*) and CXCR4 (*CXCL12*) (Figure 2F). While neutrophil chemokine expressing FB3 was increased in PP, the neutrophil chemoattracts that bind CXCR2 were not well detected in this condition. Taken together, these results demonstrate that SS is associated with a prominent IFN and suggest that fibroblast recognition of IFN may drive neutrophil recruitment in SS.

We next interrogated the lymphoid, myeloid, and keratinocyte subsets in our data to understand whether other cell types respond to IFN in SS (Figure S2A-C). 9 transcriptionally distinct lymphocyte subpopulations were identified by dimensionality reduction and unsupervised clustering of the initial lymphoid and plasma cell clusters (Figure S2A). Decreased in frequency across neutrophilic dermatoses compared to HC were stem-like CD4 T cells LYM1 (*IL7R, TCF7*), CD8+ T cells LYM4 (*CD8A*), and NK cells LYM7 (*KLRD1*) (Figure S2D). B cells and plasma cells, LYM2 (*IGLC3, IGHG2, IGKC*) and LYM9 (*IGHM, IGLC1*), were enriched in PG and PP. Activated CD4 T cells LYM5 (*CD69*) were increased in frequency in PG but not PP or SS. IFN-activated lymphocytes LYM8 (*MX1, OAS3, IFI44L*) were found predominantly in SS.

We next investigated keratinocyte subsets and found by dimensionality reduction and unsupervised clustering 9 transcriptionally discrete clusters (Figure S2B). Basal keratinocytes KC1 (*KRT15*) and suprabasal keratinocytes KC3 (*KRT1*, *KRT10*) were decreased in frequency across neutrophilic dermatoses, whereas stratum granulosum keratinocytes KC2 (*KLK13*) and IFN-activated keratinocytes KC8 (*IRF1, WARS, CXCL10*) were increased (Figure S2F). IFN-activated KC8 was observed at 3-fold higher frequency in SS compared with other neutrophilic dermatoses.

Dimensionality reduction and unsupervised clustering of the initial myeloid, pDC, DC, and mast cell clusters resolved 10 transcriptionally distinct myeloid cell clusters (Figure S2C). MY4 mast cells (*GATA2, HDC, TPSAB1*) and MY3 macrophages *(SELENOP, APOE, RNASE1)* were enriched in HC skin, whereas MY6 DCs (*CCR7, CCL22*), and MY7 pDCs (*GZMB*) were increased across neutrophilic dermatoses (Figure S2F). TREM2 macrophages MY5 (*SPP1*) were found in SS and PG but not PP nor HC skin. IFN-activated MY1 (*CXCL10, GBP5, CD300E*) were enriched in SS and PG but not PP.

Pathway analysis indicated that LYM8, KC8, and MY1 may be regulated by IFN response factor 1 (IRF1) and STAT1, confirming the identity of these subsets as IFN-activated (Figure S2G-I). Thus, IFN-activated subsets were expanded in all major skin cell lineages in SS skin.

We next determined the IFN signature in individual SS patients (Figure S3A-E). Each SS patient sample possessed a unique composition of IFN-activated cell types. IFN-activated myeloid cells were observed in all SS patients to a varying degree, whereas IFN-activated myeloid, lymphocyte, and keratinocyte were observed in a subset of patients. These results demonstrate that the prominent IFN signature is a conserved feature of SS but that the cell type on which IFN acts is patient dependent.

To validate our snRNA-Seq findings, we next performed bulk RNA-Seq on a large independent cohort of SS (n=45), PG (n=58), and HC skin (n=4) (Figure 3A). Cell type deconvolution confirmed snRNA-Seq findings of increased frequency of DCs, lymphoid cells, myeloid cells, pDCs, plasma cells, and proliferating cells in neutrophilic dermatoses (Figure 3B). Differential expression analysis identified genes upregulated in both SS and PG including proinflammatory cytokines (*IL1A, IL1B, TNF, IL6, OSM*), neutrophil growth factor (*CSF3*), antimicrobials (*LCN2, CAMP*), neutrophil chemokine receptors CXCR2 and CXCR4, neutrophil chemokines that bind CXCR2 (*CXCL1, CXCL2, CXCL3, CXCL5, CXCL6, CXCL8*) (Figure S4A-B). Overall, the differentially expressed genes shared between SS and PG were related to inflammatory pathways including ‘innate immune response’ and ‘neutrophil degranulation’ (Figure S4C).

**Figure 3:**
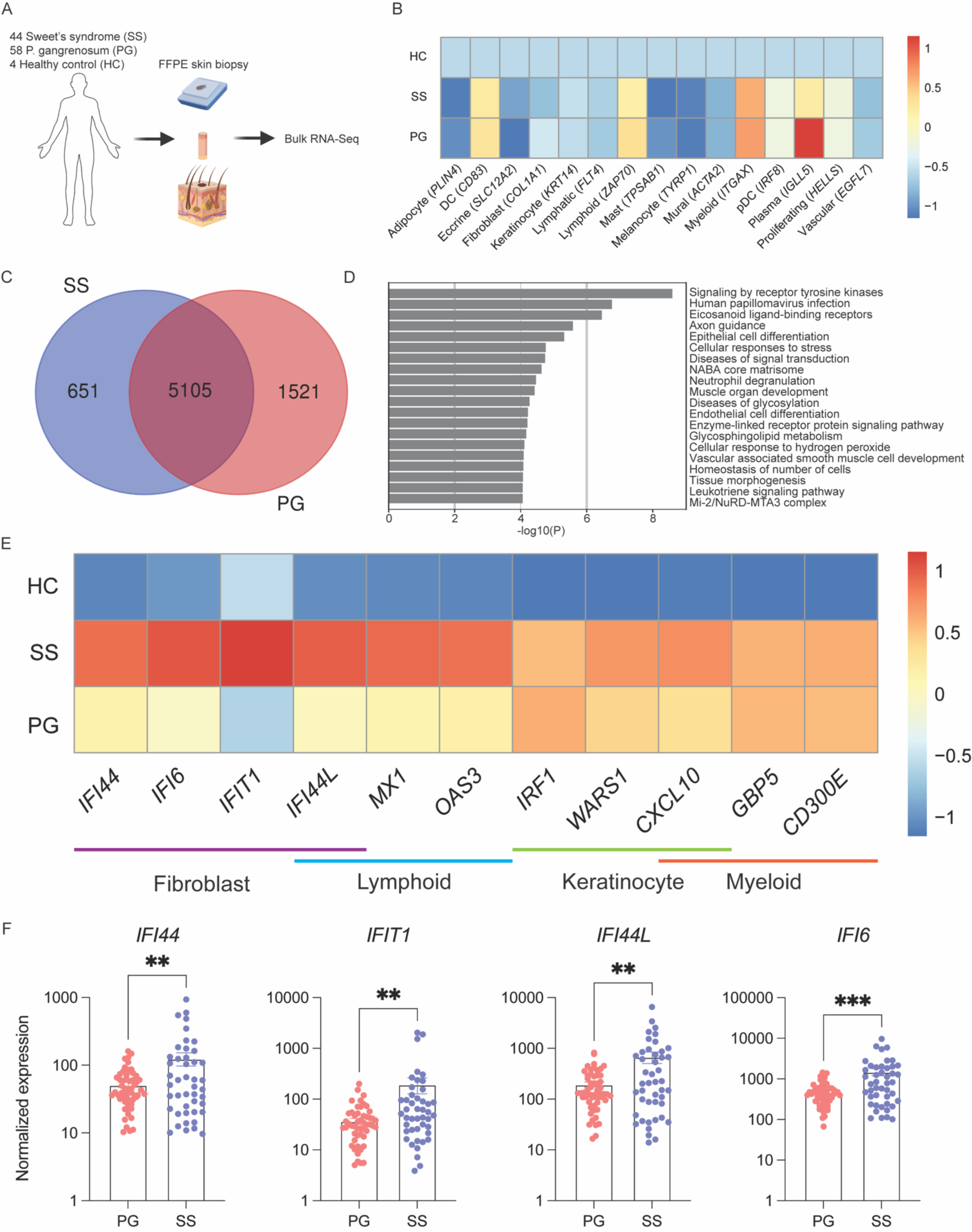
Bulk RNA-Seq of neutrophilic dermatoses validates prominent IFN signature in SS. (A) Experimental schematic. (B) Cell type deconvolution of bulk RNA-Seq using marker genes from snRNA-Seq. (C) Identification of differentially expressed genes (up and down, P_adj_ < 0.05) unique to SS compared to HC and PP. (D) Pathway analysis with the 651 differentially expressed genes unique to SS. (E) Bulk RNA-Seq normalized gene expression of IFN-induced genes marking each cellular compartment in snRNA-Seq data, scaled by column. (F) Bulk RNA-Seq normalized gene expression of IFN-induced genes marking IFN-activated FB6. Each dot represents one patient. **P < 0.01 and ***P < 0.001 using unpaired t test. SS, Sweet’s syndrome; PG, pyoderma gangrenosum; HC, healthy control; IFN, interferon; FB, fibroblast.

Comparative analysis identified 651 and 1521 genes uniquely regulated in SS and PG, respectively, and 5105 similarly regulated genes (Figure 3C). Proinflammatory cytokines *IL1A*, *IL1B*, *TNF*, and *IL-6* and neutrophil chemokines *CXCL1, CXCL2, CXCL3, CXCL5, CXCL6*, and *CXCL8* were significantly upregulated in both SS and PG. The genes upregulated in both conditions were consistent with ‘innate immune response’ and ‘neutrophil degranulation’ (Figure S4C). *IL17A* was not significantly upregulated in PG or SS (Figure S4-B). Pathway analysis suggested PG-specific genes—including *COL11A1*—were consistent with developmental processes and extracellular matrix production, validating our snRNA-Seq findings (Figure S4D). SS-specific genes were consistent with ‘Human papilloma virus infection’—a type I IFN response pathway (Figure 3D). Expression of the 11 IFN stimulated genes defining IFN-activated subsets of fibroblasts, myeloid cells, lymphoid cells, and keratinocytes was significantly increased in SS compared to HC (Figure 3E). 10 of these genes—particularly those marking IFN-activated fibroblasts and lymphocytes—were increased in SS over PG. This result supports our snRNA-Seq observation of IFN-activated fibroblast and lymphocyte subsets uniquely present in SS lesions. Genes marking IFN-activated fibroblasts *IFI44, IFI6, IFI44L,* and *IFIT1* were highly expressed in many SS patients (Figure 3F). Taken together, this unbiased analysis of a large independent patient cohort validates the prominent IFN signature identified in SS by snRNA-Seq.

### Type I interferon activates primary human dermal fibroblasts to highly express neutrophil chemokines

To test the significance of the IFN signature in fibroblasts for neutrophil recruitment, we cultured primary human dermal fibroblasts from 50 different donors with type I IFN (IFNa2) and type 2 IFN (IFNγ) and examined gene expression by bulk RNA-Seq (Figure 4A). The effect of these signals was compared with TNFα, which has been shown to induce fibroblast expression of neutrophil chemokines (Cavagnero et al., 2024, Cavagnero and Gallo, 2022). Comparative differential expression analysis identified 76 genes highly upregulated by IFNγ and IFNa2 but not TNFα (Figure 4B). Among these genes were the IFN stimulated genes that marked IFN-activated snRNA-Seq FB6 (Figure 4C). Highly upregulated by all 3 stimuli were chemoattractants for monocytes/macrophages (*CCL2, CCL7, CCL8, CX3CL1*) and lymphocytes (*CXCL9, CXCL10, CXCL11*) (Figure S5A). Few inflammatory genes were highly upregulated by IFNγ alone (*CCL13*) or by TNFα alone (*IL34, TSLP*) (Figure S5A). Remarkably, the 85 genes highly upregulated by TNFα and IFNa2 but not IFNγ included neutrophil chemokines (*CXCL1*, *CXCL5*, *CXCL8, C3)*, myeloid cell growth factors (*CSF2*, *CSF3*), and proinflammatory cytokines associated with SS (*IL1A, IL1B, IL6, TNF*) (Figure 4D). Neutrophil chemokine upregulation by type I IFN was validated by qPCR (Figure 4E, S5B).

**Figure 4:**
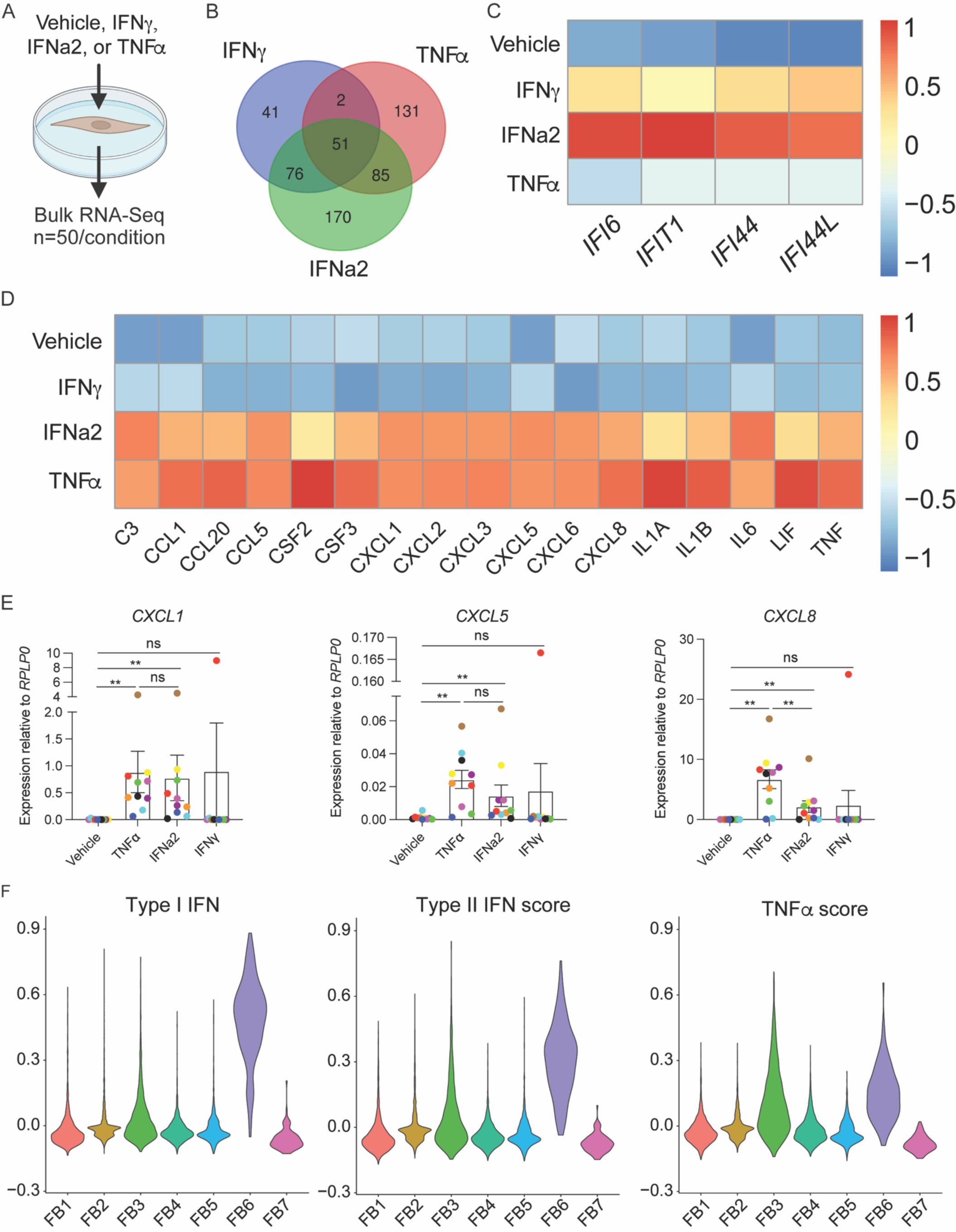
Cultured primary human dermal fibroblasts are activated by type 1 IFN to express neutrophil chemokines. (A) Schematic of *in vitro* assay to test significance of IFN signature in primary human dermal fibroblasts. (B) Venn diagram comparing the number of genes highly upregulated (P_adj_ < 0.05, Log_2_ fold change > 3) per stimuli. (C) Bulk RNA-Seq normalized gene expression of IFN-induced genes marking FB6 in stimulated fibroblasts that were highly upregulated by IFNγ and IFNa2 but not TNFα. (D) Bulk RNA-Seq normalized gene expression of select neutrophil chemokines, myeloid growth factors, and proinflammatory cytokines that were highly upregulated by TNFα and IFNa2 but not IFNγ. (E) qPCR normalized gene expression of neutrophil chemokines in stimulated dermal fibroblasts. Each color represents a single donor’s fibroblasts. ns (not significant), *P < 0.05, **P < 0.01, ***P < 0.001, and ****P < 0.0001 using Wilcoxon matched-pairs signed rank test. (F) IFNγ, IFNa2, and TNFα signature scores in snRNA-Seq fibroblast subsets using all significantly upregulated genes (P_adj_ < 0.05) by each cytokine *in vitro*. SS, Sweet’s syndrome; PG, pyoderma gangrenosum; PP, pustular psoriasis; HC, healthy control; IFN, interferon.

To infer whether type I or type II IFN drove the IFN signature in FB6, *in vitro* activation gene modules were generated and projected onto snRNA-Seq data. IFN-activated FB6 demonstrated a transcriptomic signature most consistent with activation by type I IFN (Figure 4F). Together, these findings support a model wherein type I IFN activates subsets of fibroblasts in SS to recruit neutrophils.

### Spatially distinct fibroblast subsets promote neutrophil recruitment in Sweet’s syndrome

We next sought to understand the cellular and molecular landscape of neutrophilic dermatoses biogeographically and performed subcellular resolution spatial transcriptomics of archival FFPE skin biopsies from SS (n=5), PP (n=2), and HC (n=2) using Nanostring’s 1000 gene imaging-based CosMx platform (Figure 5A). At the single-cell level, our dataset included 39992 cells, with an average of 54 genes and 136 transcripts detected per cell. Dimensionality reduction and unsupervised clustering revealed 10 transcriptionally distinct clusters (Figure 5B-C): neutrophils (*CCL3L3, CXCL8, IL1B*), keratinocyte cluster 1 (*KRT6C, KRT6B, KRT16*), keratinocyte cluster 2 (*KRT5, KRT14, S100A2*), fibroblasts (*COL6A2, COL3A1, COL1A1*), myeloid cells (*HLA-DPA1, HLA-DRB1, CD74*), lymphoid cells (*MALAT1, CCL19, IL32*), mural cells (*VIM, COL4A2, IGFBP7*), keratinocyte cluster 3 (*TNFSF15, KRT1, KRT10*), and keratinocyte cluster 4 (*IL36G, S100A8, S100A9*). The frequencies of neutrophils, myeloid cells, and lymphoid cells were increased in neutrophilic dermatoses compared to HC (Figure 5D), confirming snRNA-Seq and bulk RNA-Seq findings.

**Figure 5:**
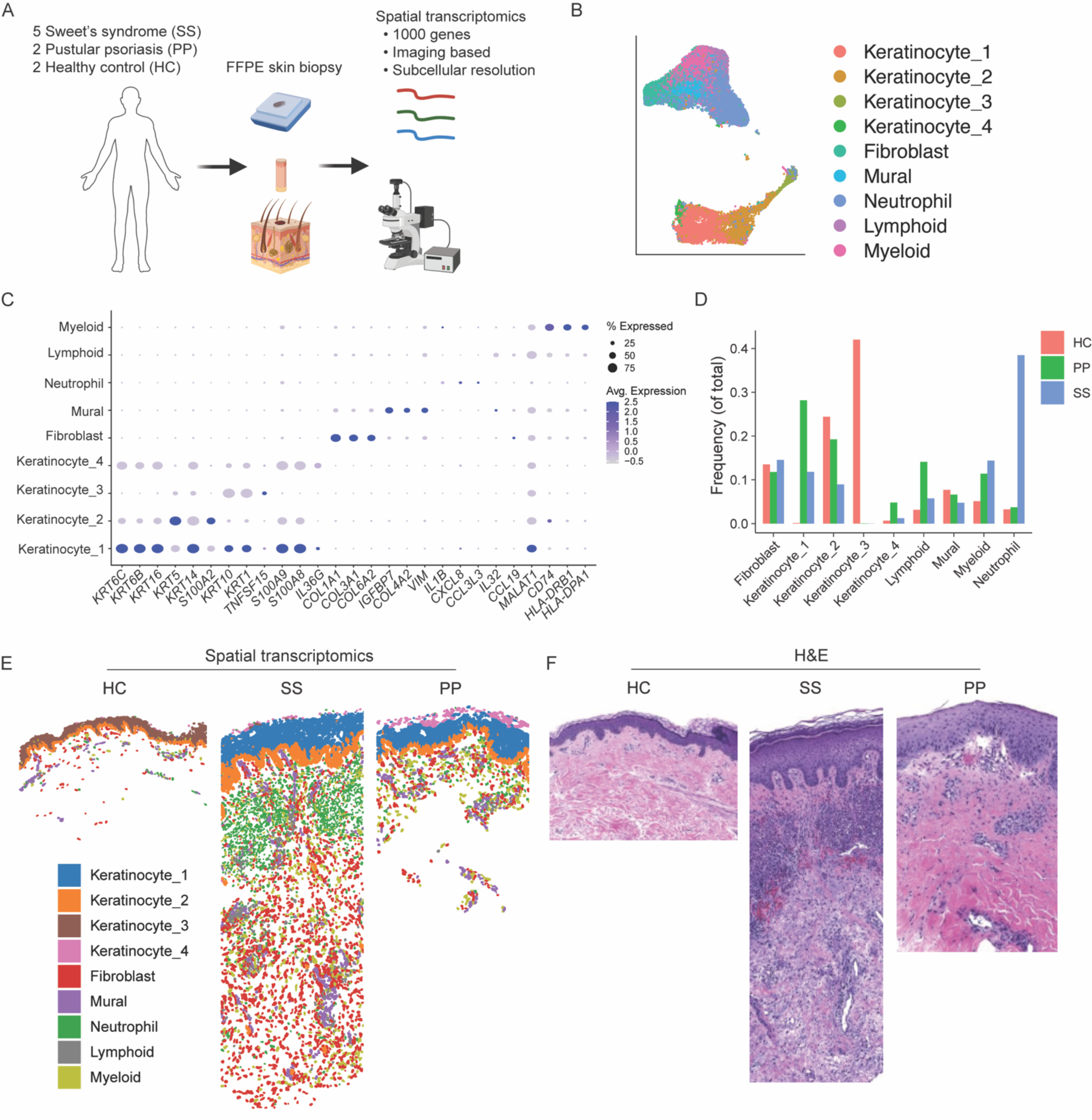
Single-cell resolution spatial transcriptomics of neutrophilic dermatoses. (A) Experimental schematic using the Nanostring CosMx subcellular resolution spatial transcriptomics platform. (B) Dimensionality reduction and unsupervised clustering, colored by cell type. (C) Expression of top 3 marker genes per cluster. (D) Proportion of each cluster across conditions. (E) Clusters projected onto representative tissue sections from HC, SS, and PP. (F) H&E staining of adjacent tissue sections from HC, SS, and PP. SS, Sweet’s syndrome; PP, pustular psoriasis; HC, healthy control; H&E, hematoxylin and eosin.

Cell clusters were then projected onto tissue sections to elucidate spatial localization (Figure 5E). Strikingly, all unsupervised clusters mapped to unique histological regions. SS and PG shared with HC basal epidermis keratinocyte cluster 2 but had distinct spinous and supraspinous epidermal clusters. The supra-basal epidermal clusters in SS and PP highly expressed proinflammatory genes (*IL36, S100A8, S100A9*) whereas those in HC did not. Unlike single-cell and single-nucleus transcriptomics, which often fail to recover neutrophils, spatial transcriptomics identified a cluster of neutrophils that was validated by H&E staining (Figure 5F). Neutrophils in SS were observed densely packed in the mid-to-upper dermis, whereas neutrophils in PP were more evenly dispersed throughout the upper dermis. Differential expression analysis of neutrophils across conditions identified upregulation of *IL1B*, *CXCL8*, and *IL1RN* in SS neutrophils, supporting previous work demonstrating overexuberant neutrophil activation in SS (Figure S6) (Bhattacharya et al., 2023).

We next performed unsupervised niche analysis by clustering cells into 7 regions based on the cellular composition within a 50um radius of each cell using MClust (Scrucca et al). This analysis resolved an SS associated upper dermal niche (niche1) and an SS associated lower dermal niche (niche2) (Figure 6A-B, S7A-B). Niche1 included fibroblasts and neutrophils, whereas niche2 included fibroblasts but not neutrophils (Figure 6C). Differential expression analysis between fibroblasts in niche1 and niche2 indicated that fibroblasts in the upper dermal, neutrophil proximal niche were marked by IFN inducible neutrophil chemokine *CXCL5*, whereas fibroblasts in the lower dermal, neutrophil distal niche were marked by IFN inducible gene *IFITM3*, neutrophil chemokines *CXCL12* and *MIF*, and extracellular matrix genes (*COL3A1, COL1A1, COL1A2*) (Figure 6D). Similar results were obtained both by performing differential expression analysis between neutrophil proximal (<10um) and distal (>50um) fibroblasts and by correlating fibroblast gene expression with neutrophil distance (Figure 6E-G). Projection of *IFITM3*, *CXCL1*, and *CXCL12* expression onto tissue sections indicated that these genes were not expressed in HC skin and highly expressed in neutrophilic dermatoses lesions (Figure 6H-J, S6C-H). *IFITM3* demonstrated striking regional expression in the epidermis with high expression in basal but not suprabasal keratinocytes. Protein immunostaining confirmed that CXCL1 was expressed by a greater frequency of fibroblastic cells in the upper dermis, whereas CXCL12 was expressed by a greater frequency of fibroblastic cells in the lower dermis (Figure 6K). Thus, unbiased subcellular resolution spatial transcriptomics identified positionally distinct dermal immune acting fibroblast subsets in SS lesions.

**Figure 6:**
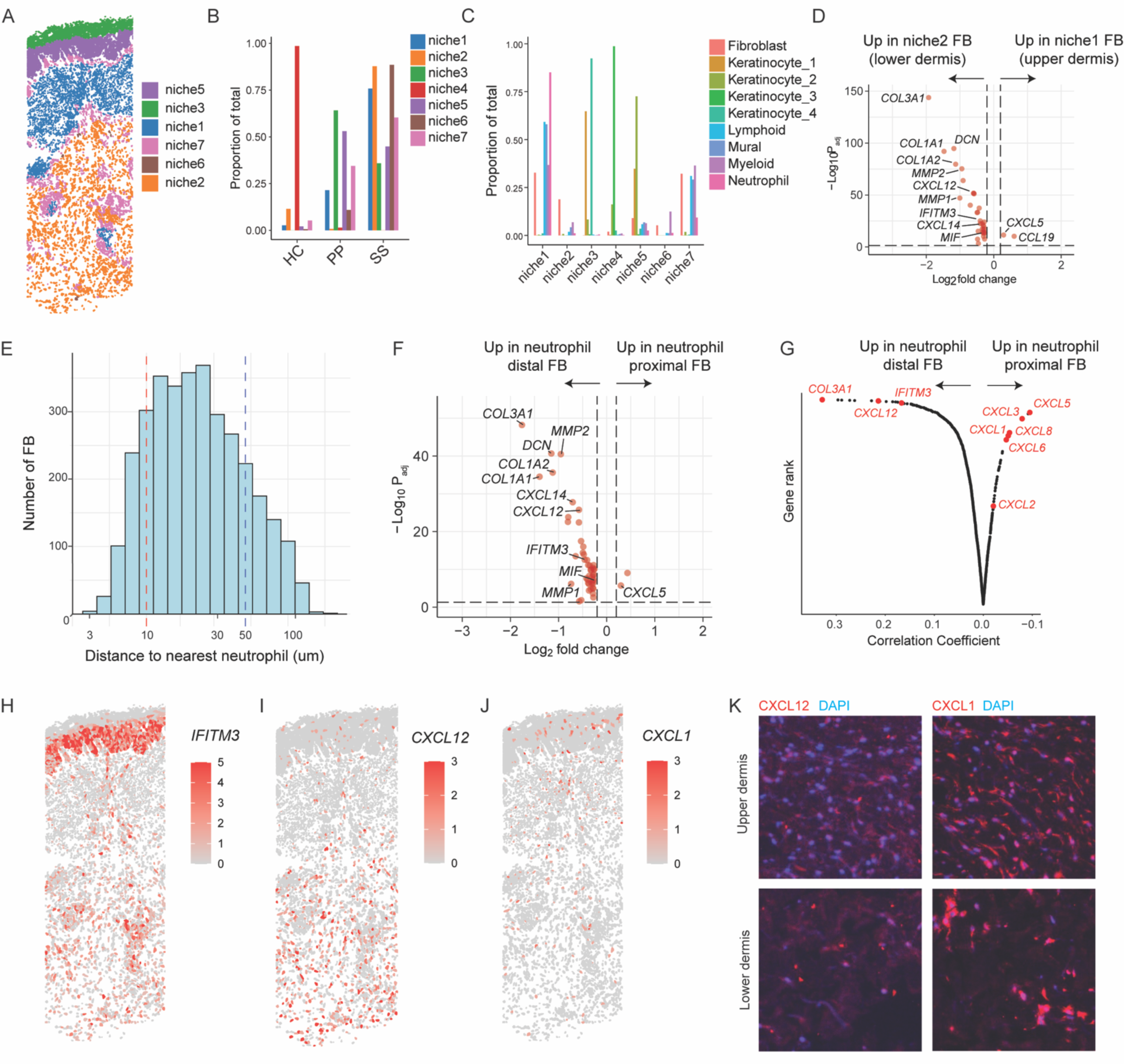
Niche analysis reveals positionally distinct immune acting fibroblasts in SS. (A) Niche analysis projected onto SS tissue. (B) Frequency of each niche per condition. (C) Frequency of each cell type per niche (includes all conditions). (D) Differential expression analysis between fibroblasts in upper dermal niche1 and fibroblasts in lower dermal niche2. Adjusted (adj) P value threshold = 0.05. Log_2_ fold-change (FC) threshold = 0.2. (E) Distribution of fibroblast distance to nearest neutrophil. (F) Differential expression between neutrophil proximal fibroblasts (<30μm) and neutrophil distal fibroblasts (>300μm). Adjusted (adj) P value threshold = 0.05. Log_2_ fold-change (FC) threshold = 0.2. (G) Rank order of genes based on correlation with fibroblast distance to nearest neutrophil. SS expression of interferon-induced gene *IFITM3* (H) and neutrophil chemokines *CXCL12* (I) and *CXCL1* (J). (K) SS upper dermal and lower dermal immunostaining of CXCL1 (left) and CXCL12 (right). SS, Sweet’s syndrome.

## Discussion

In this study, we aimed to develop a comprehensive cellular and molecular atlas of the human neutrophilic dermatosis Sweet’s syndrome (SS)—a rare disease not previously profiled at scale using unbiased transcriptomics. Comparative analysis of single-nucleus transcriptomics from archival clinical skin biopsies of patients with SS, early pyoderma gangrenosum (PG), pustular psoriasis (PP), and healthy controls (HC) revealed new cellular contributors to neutrophilic dermatoses, including pDCs. This analysis also led to the identification of an IFN-activated fibroblast subset in SS lesions that was absent in other conditions. Bulk RNA-Seq of a large, independent patient cohort confirmed the snRNA-Seq findings, including a unique and prominent IFN signature in SS. Subsequent unbiased functional experiments demonstrated that cultured primary human dermal fibroblasts—cells mainly appreciated for supporting tissue architecture—highly expressed neutrophil chemokines that bind CXCR2 following recognition of type I IFN. These results are consistent with a 2024 report showing that type I IFN induced CXCL8 secretion in hepatocellular carcinoma cells (Ma et al., 2024).

Fibroblasts have historically been regarded as a homogenous cell type that supports tissue architecture but are becoming appreciated as a diverse and multifunctional class of cells (Cavagnero and Gallo, 2022). Subsets of immune acting fibroblasts (IAFs) play a critical role in IL-17-mediated neutrophil recruitment during *S. aureus* infection and psoriasis vulgaris through secretion of chemokines that bind CXCR2 and CXCR4 (Cavagnero et al., 2024). Unlike psoriasis vulgaris, biologics targeting the IL-17 pathway have not demonstrated efficacy in treating SS (de Risi-Pugliese et al., 2019). Therefore, we hypothesized that IAFs drive neutrophil recruitment in SS through an IL-17 independent mechanism. We now present evidence that human dermal IAFs play a role in neutrophil recruitment in SS through recognition of type I IFN.

A 2022 study identified type I IFN-activated dermal fibroblasts in the autoimmune skin disease cutaneous lupus erythematosus using scRNA-Seq (Billi et al., 2022). While type I IFN and neutrophils are well known to drive lupus pathogenesis (Bruera et al., 2022), the connection between type I IFN and neutrophil recruitment has not been previously established. In light of our study—the first to present data supporting a role for type I IFN-activated dermal fibroblast subsets in neutrophil inflammation—these findings suggest that IAF recognition of type I IFN may contribute to neutrophil inflammation in other autoimmune diseases and interferonopathies.

Psoriasis vulgaris skin cell atlas studies have resolved IAF subsets that express neutrophil chemokines that bind CXCR2 and CXCR4 (Ma et al., 2023, Cavagnero et al., 2024). Here, using a combination of subcellular resolution spatial transcriptomics and immunostaining, we identified two distinct neutrophil chemokine-expressing fibroblast populations: a neutrophil-proximal subset expressing CXCR2 ligands in the upper dermis, and a neutrophil-distal subset expressing the CXCR4 ligand CXCL12 in the lower dermis. These findings suggest that upper and lower dermal fibroblasts are capable of adopting a neutrophil recruiting IAF state depending on the disease context.

This current study also identified subsets of IFN-activated keratinocytes, myeloid cells, and lymphocytes enriched in SS. Similar subpopulations have been reported in cutaneous lupus erythematosus (Billi et al., 2022), hidradenitis suppurativa (van Straalen et al., 2024, Gudjonsson et al., 2020), and psoriasis (Ma et al., 2023). Future research should focus on understanding how IFN action on these cell types contributes to SS pathogenesis.

Given the diversity of conditions associated with SS, some speculate that it may lack a common underlying molecular mechanism (Joshi et al., 2022). However, type I IFN has been implicated in these associated conditions, including inflammatory disease, infection, vaccination, pregnancy, and cancer. Furthermore, numerous case studies have documented SS induced by IFN therapy (Rodriguez-Lojo et al., 2014, Gheorghe et al., 2008, Kim et al., 2015). Thus, type I IFN may serve as a unifying factor in SS, suggesting that FDA approved JAK/STAT inhibitors, which block type I IFN signaling, could offer a new therapeutic strategy for SS. Indeed, a 2020 case study reported significant improvement of SS with baricitinib treatment (Nousari et al., 2021).

Current models of SS pathogenesis—developed from limited laboratory studies of serum and skin inflammatory mediators and case reports of targeted therapeutic trials—are incomplete but suggest that infection, cancer, and drug reactions drive heightened levels of TNFα, IL-1, and G-CSF that promote leukocytosis (Heath and Ortega-Loayza, 2019). In the context of hematological cancer, aberrant malignant neutrophils further contribute to leukocytosis following treatment with G-CSF, all-trans retinoic acid, or FL3 inhibitor. Th1, Th2, and/or Th17 cells are thought to then promote IL-17, TNFα, and/or IL-1 dependent neutrophil recruitment, activation, and neutrophil extracellular trap (NET) formation (Joshi et al., 2022). Genetic variants may increase the risk of developing SS independent of malignancy. For example, a 2023 case study of a patient with non-cancer associated SS identified a gain-of-function *PIK3R1* mutation—specifically in neutrophils—that increased neutrophil migration and respiratory burst capacity (Bhattacharya et al., 2023).

Consistent with current models, we observed increased expression of proinflammatory cytokines, neutrophil growth factors, and neutrophil chemokines in SS compared to healthy skin including IL-1, TNFα, G-CSF (encoded by *CSF3*), and CXCR2 ligands. Further, spatial transcriptomics enabled in situ characterization of SS neutrophils, revealing that these cells expressed more IL-1β and CXCL8 than neutrophils in both healthy and PG skin. Contrary to the current model, but consistent with a recent case report (de Risi-Pugliese et al., 2019), our data do not support a role for IL-17 in SS pathogenesis.

Based on our findings of increased pDCs in SS, a prominent and unique IFN signature, type I IFN activation of dermal fibroblasts to express neutrophil chemokines, and upper dermal localization of fibroblast subsets expressing CXCR2 binding neutrophil chemokines, we speculate that inciting factors drive the production of type I IFN, IL-1, and TNFα. These proinflammatory cytokines then promote subsets of dermal fibroblasts to recruit neutrophils to infiltrate the skin and release extracellular traps, which in turn activate pDCs to secrete type I IFN, creating a self-sustaining inflammatory positive feedback loop.

Our study is limited in that our snRNA-Seq cohort included a single PP sample. Because our aim was to define SS pathogenesis, not necessarily PP and PG, it is important that this PP sample be included as it adds value as a comparator. Importantly, however, conclusions about the PP landscape should not be made based off this single patient biopsy. Nevertheless, our spatial transcriptomics data included 2 additional PP patient samples that aligned well with the PP snRNA-Seq data.

In summary, we have used cutting-edge single nucleus, bulk, and spatial transcriptomics on archival clinical samples to illuminate the pathogenesis of rare human neutrophilic dermatoses. These approaches led to the identification of a unique IFN signature in SS, which was seen prominently in a subset of upper dermal fibroblasts that highly express neutrophil chemokines following recognition of type I IFN. This work provides insight into the mechanisms underlying the clinical observation of recombinant type I IFN-induced SS and identifies type I IFN and IFN-activated fibroblasts as a novel therapeutic target for the treatment of SS. The comprehensive cell atlas generated here is anticipated to be a valuable resource for the research community to facilitate future discoveries.

## Methods

Single-nucleus RNA sequencing: FFPE samples of skin biopsies were obtained from patients with SS (n=6), PP (n=1), PG (n=6), and HC (n=5). Libraries were generated using 10X Genomics Flex FFPE protocol and were subjected to 28×91bp of sequencing according to the manufacturer’s protocol (Illumina NovaSeq). Library prep and next-generation sequencing was carried out in the Advanced Genomics Core at the University of Michigan. CellRanger with default parameters was used to perform alignment to the hg38 reference genome and gene counting. Data were filtered, processed, and analyzed using Seurat (Butler et al., 2018; Stuart et al., 2019). All functions described below are Seurat functions unless stated otherwise. Filtering data involved removing low quality cells, removing ambient RNA with SoupX with setContaminationFraction = 0.2 (Young and Behjati, 2020), and removing doublets using DoubletFinder with the default settings (McGinnis et al., 2019). Data were normalized and integrated using NormalizeData and IntegrateData with default parameters. Clusters were identified using FindNeighbors with 50 principal components and FindClusters with a range of resolutions. For each resolution, nonlinear dimensionality reduction and visualization was performed with 50 principal components, and marker genes for each cluster were determined using FindAllMarkers with min.pct = 0.25. The resolution yielding clusters with the most distinct marker genes was chosen for further analysis. For cell subpopulation analysis, data were subset based on cell type annotation, contaminating cells were removed, and the above analysis was repeated. Pathway analysis was performed with Metascape (Zhou et al., 2019). Signature scores were generated using AddModuleScore with all significantly upregulated genes (P_adj_ < 0.05) from *in vitro* bulk RNA-Seq.

Spatial transcriptomics: FFPE skin biopsies were obtained from patients with SS (5), PP (2), and HC (2). Subcellular resolution spatial transcriptomics was performed using the NanoString CosMx SMI platform with the predefined human 1000 gene panel as previously described (He et al., 2022). DAPI staining and immunofluorescent staining of PanCK, CD298/B2M, CD45, and SMA facilitated cell segmentation with machine learning algorithm Cellpose (Stringer et al., 2021). Counts were assigned to individual cells based on cell segmentation borders. Cells with fewer than 20 counts or with area more than 5 times the average cell area were removed. Data were normalized to total counts per cell and square root transformed. UMAP dimensionality reduction was then performed using all genes. Unsupervised Leiden clustering was run with 50 principal components and a resolution of 0.4. The following analysis was performed using Seurat (Butler et al., 2018; Stuart et al., 2019) unless otherwise noted. Differential expression using FindAllMarkers with min.pct = 0.25 was used to identify cluster markers genes. Clusters were annotated manually based on known marker genes. For niche analysis, the cellular composition within a 50um radius of each cell was determined and clustered into 7 niches using MClust (Scrucca et al., 2016). Differential expression between fibroblasts based on niche or distance to neutrophils was performed using FindMarkers with min.pct = 0.25.

Cell culture: Healthy human dermal fibroblasts were isolated from 2x 4mm skin punches by mechanical digestion with scissors and enzymatic digestion with 0.2% collagenase for 30’ at 37C. Cells were grown in a humidified incubator at 5% CO2 and 37°C under sterile conditions and used at passage 3. Cells were grown in RPMI supplemented with L-glutamine, 10% FBS, and antibiotic-antimycotic (ThermoFisher Scientific, #15240062). Cells at 80% confluency were stimulated with recombinant cytokines including rhIFNa2 (R&D, #11100-1, 5ng/ml), rhTNFα (R&D, #210-TA-005, 10ng/ml), or rhIFNγ (R&D, #285-IF-100, 5ng/ml). After 6 hours of stimulation, cells were lysed with RNA lysis buffer (Qiagen, 74104).

RNA isolation protocol: For bulk RNA-Seq and qPCR of FFPE tissue, RNA was isolated from FFPE tissue using the RNeasy DSP FFPE Kit (Qiagen, #73604). For bulk RNA-Seq and qPCR of cultured fibroblasts, RNA was isolated from cell lysates using the RNeasy kit (Qiagen, #74104).

Bulk RNA-Seq: Libraries were generated using a QuantSeq 3’ mRNA-Seq Library Prep Kit (Lexogen) and sequenced using a NovaSeq (Illumina). After adaptor trimming, reads were mapped to the hg38 reference genome using STAR, count matrices were generated using HTSeq, and differential expression analysis was performed using DESeq2. Volcano plots and heatmaps were generated using EnhancedVolcano and pheatmap. Pathway analysis was performed with Metascape (Zhou et al., 2019).

RT-qPCR: RNA was converted to cDNA using a High-Capacity cDNA Reverse Transcription Kit (ThermoFisher, # 4368814). qPCR was performed using a QuantStudio Real-Time PCR system (ThermoFisher) with TaqMan Universal PCR Master Mix (ThermoFisher, #4304437). Housekeeping gene *RPLP0* was used to normalize expression. TaqMan primers included *RPLP0* (ThermoFisher, #Hs004200895_gh), *CXCL1* (ThermoFisher, #Hs00236937_m1), *CXCL2* (ThermoFisher, #Hs00234140_m1), *CXCL3* (ThermoFisher, # Hs00171061_m1), *CXCL5* (ThermoFisher, #Hs00982282_m1), *CXCL6* (ThermoFisher, #Hs00237017_m1), *CXCL8* (ThermoFisher, #Hs00174103_m1).

Immunofluorescence: Antigen retrieval of FFPE sections was performed using Target Retrieval Solution (Dako, #S2369) as per manufacturer recommendations. Sections were blocked with serum from secondary antibody host, stained with primary antibodies overnight at 4°C, secondary antibodies for 1hr at RT, and nuclei were counterstained with DAPI. Epifluorescence images were taken using an EVOS5000. Brightness and contrast were adjusted slightly using ImageJ or Nikon elements software and applied equally across samples. Primary antibodies: CXCL1 (ThermoFisher, #PA586508, 1:100), CXCL12 (ThermoFisher, #14-7992-81, 1:1000). Secondary antibody: Cy3 Donkey anti-Rabbit IgG (BioLegend, #406402, 1:500).

Statistical analysis: snRNA-Seq and spatial transcriptomics differential expression analysis was performed using Seurat. Bulk RNA-Seq differential expression analysis was performed using DESeq2. Transcriptomics P values were adjusted (adj) for multiple hypothesis testing, and P_adj_ < 0.05 was considered statistically significant. qPCR statistical significance was calculated using GraphPad Prism with *P < 0.05, **P < 0.01, ***P < 0.001, and ****P < 0.0001.

Study approval: Human skin biopsies for spatial transcriptomics, H&E, and immunofluorescence were collected from the UCSD Dermatology Clinic. Sample acquisitions were approved and regulated by the UCSD Institutional Review Board (#140144). Acquisition of human skin samples for snRNA-Seq, bulk RNA-Seq, and *in vitro* fibroblast studies was approved by University of Michigan IRB. Written informed consent was obtained from all subjects.

Data availability: Genomic data presented here will be made publicly available following publication.

## Author contributions

K. Cavagnero—conceptualization, investigation, supervision, resources, software, formal analysis, visualization, writing (original draft), writing (review & editing); C. Dobry, F. Li, R. Bogle, J. Kirma, J. Fox, A. Coon, J. Albright, A. Billi, P. Harms, L. Tsoi—resources, formal analysis, investigation; J. Kahlenberg, B. Hinds, J. Gudjonsson—resources, supervision; R. Gallo—supervision, conceptualization, writing (review & editing).

## Acknowledgements

Research reported in this publication was supported by the University of Michigan Advanced Genomics Core, the UM Single Cell Spatial Analysis Program and the National Cancer Institutes of Health under Award Number P30CA046592 using the following Cancer Center Shared Resource: Single Cell and Spatial Analysis Shared Resource. Figure models were created using BioRender.com. KJC is supported by NSF GRFP2038238 and NIH T32DK007202, and RG is supported by NIH R01DK121760, NIH R01AR076082, NIHR01AI153185, U01AI152038, P50AR080594, and NIH R37AI052453. RLG is a consultant for and has equity interest in MatriSys bioscience and Sente Inc.

