## Supplemental Figures for "Positionally distinct interferon stimulated dermal immune acting fibroblasts promote neutrophil recruitment in Sweet’s syndrome"

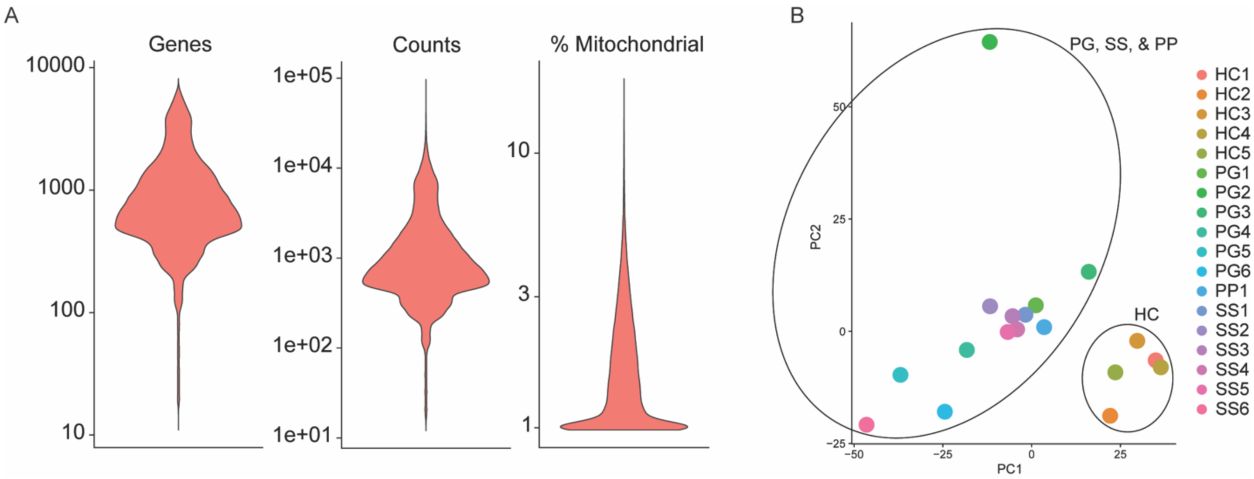


**Figure S1:** Single-nucleus RNA-Seq quality control. (A) Distribution of genes, unique counts, and percent mitochondrial reads in the snRNA-Seq dataset following removal of low-quality cells and doublets. (B) Principal component analysis of pseudo-bulked snRNA-Seq data.

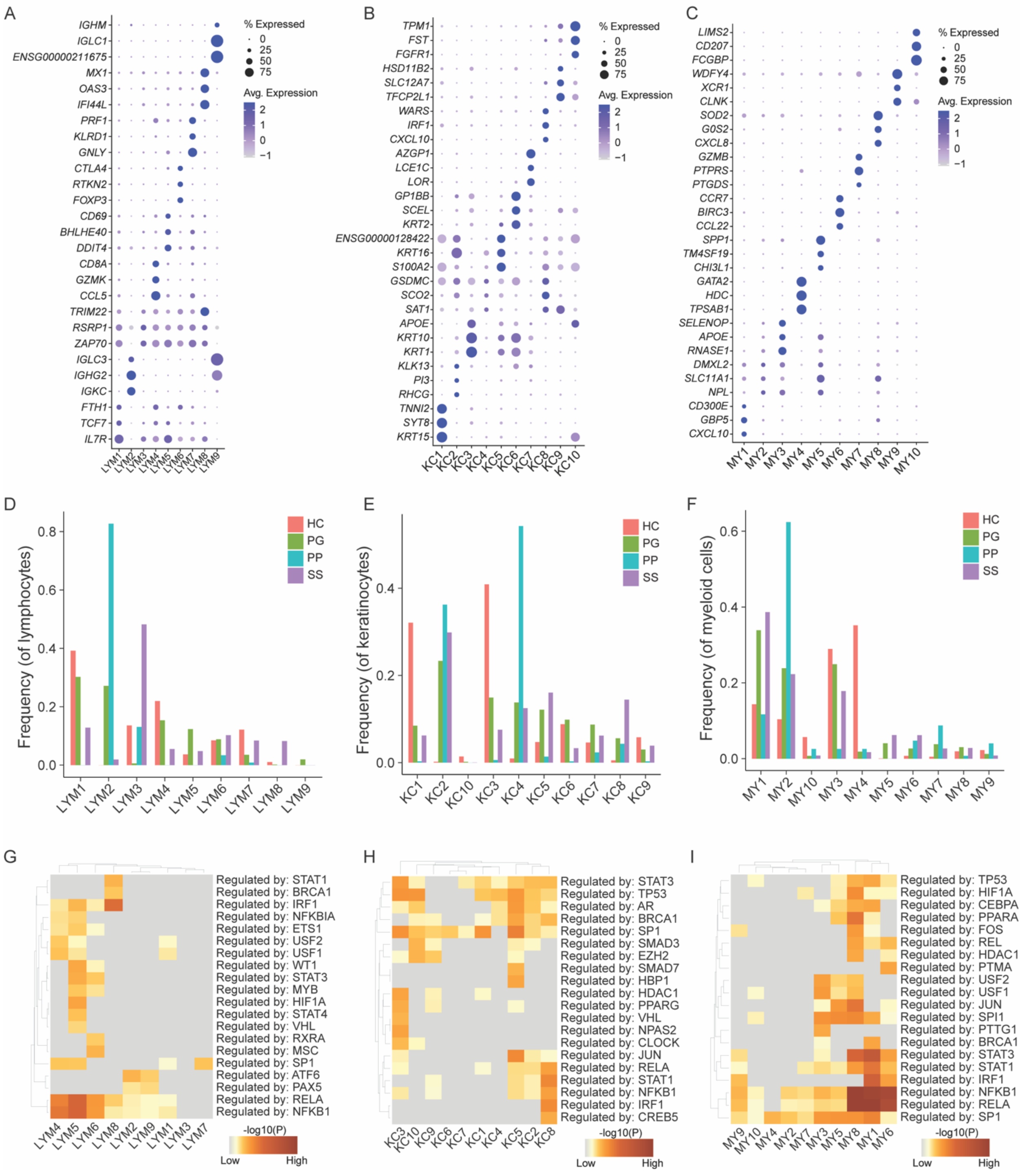


**Figure S2:** IFN-activated lymphocytes, keratinocytes, and myeloid cells are enriched in Sweet's syndrome. (A) Expression of top three marker genes for lymphocyte (LYM) cell subsets. (B) Expression of top three marker genes for keratinocyte (KC) subsets. (C) Expression of top three marker genes for myeloid cell (MY) subsets. (D) Proportion of lymphoid cell subsets across conditions. (E) Proportion of keratinocyte subsets across conditions. (F) Proportion of myeloid cell subsets across conditions. Lymphocyte (G), keratinocyte (H), and myeloid (I) subset pathway analysis using significant differentially expressed genes ($P_{adj} < 0.05$) between clusters. SS, Sweet's syndrome; PG, pyoderma gangrenosum; PP, pustular psoriasis; HC, healthy control; IFN, interferon.

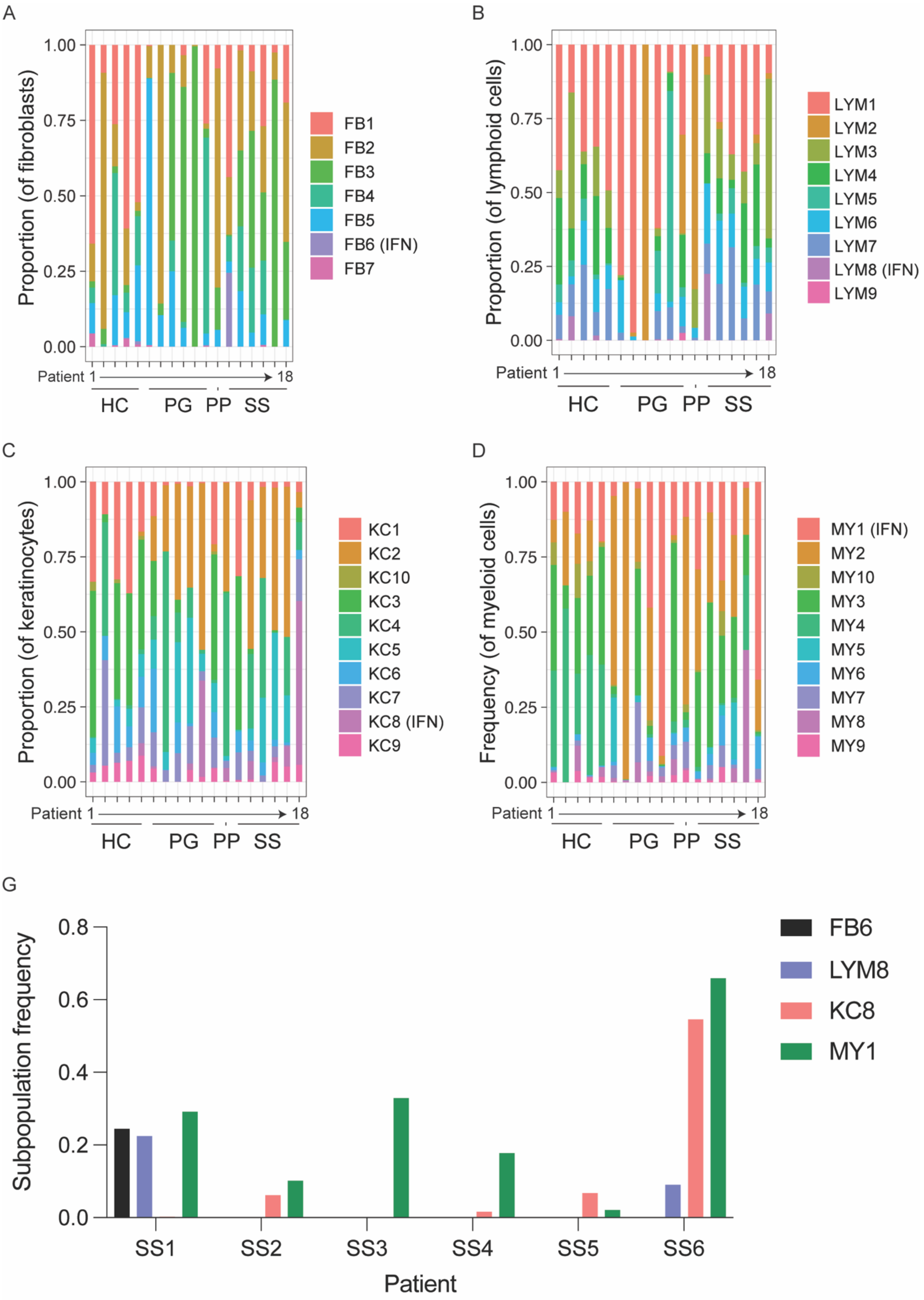

A
Proportion (of fibroblasts)
1.00
0.75
0.50
0.25
0.00
FB1
FB2
FB3
FB4
FB5
FB6 (IFN)
FB7
Patient 1
18
HC
PG
PP
SS
B
Proportion (of lymphoid cells)
LYM1
LYM2
LYM3
LYM4
LYM5
LYM6
LYM7
LYM8 (IFN)
LYM9
C
Proportion (of keratinocytes)
KC1
KC2
KC10
KC3
KC4
KC5
KC6
KC7
KC8 (IFN)
KC9
D
Frequency (of myeloid cells)
MY1 (IFN)
MY2
MY10
MY3
MY4
MY5
MY6
MY7
MY8
MY9
G
Subpopulation frequency
0.8
0.6
0.4
0.2
0.0
FB6
LYM8
KC8
MY1
SS1
SS2
SS3
SS4
SS5
SS6
Patient

**Figure S3:** Cellular compartment with IFN signature varies between Sweet's syndrome patients. (A) Proportion of fibroblast cell subsets (FB) per patient. (B) Proportion of lymphoid cell subsets (LYM) per patient. (C) Proportion of keratinocyte subsets (KC) per patient. (D) Proportion of myeloid cell subsets (MY) per patient. (E) Frequency of IFN-activated subpopulations (of parent population) split by individual SS patient. SS, Sweet's syndrome; PG, pyoderma gangrenosum; PP, pustular psoriasis; HC, healthy control; IFN, interferon.

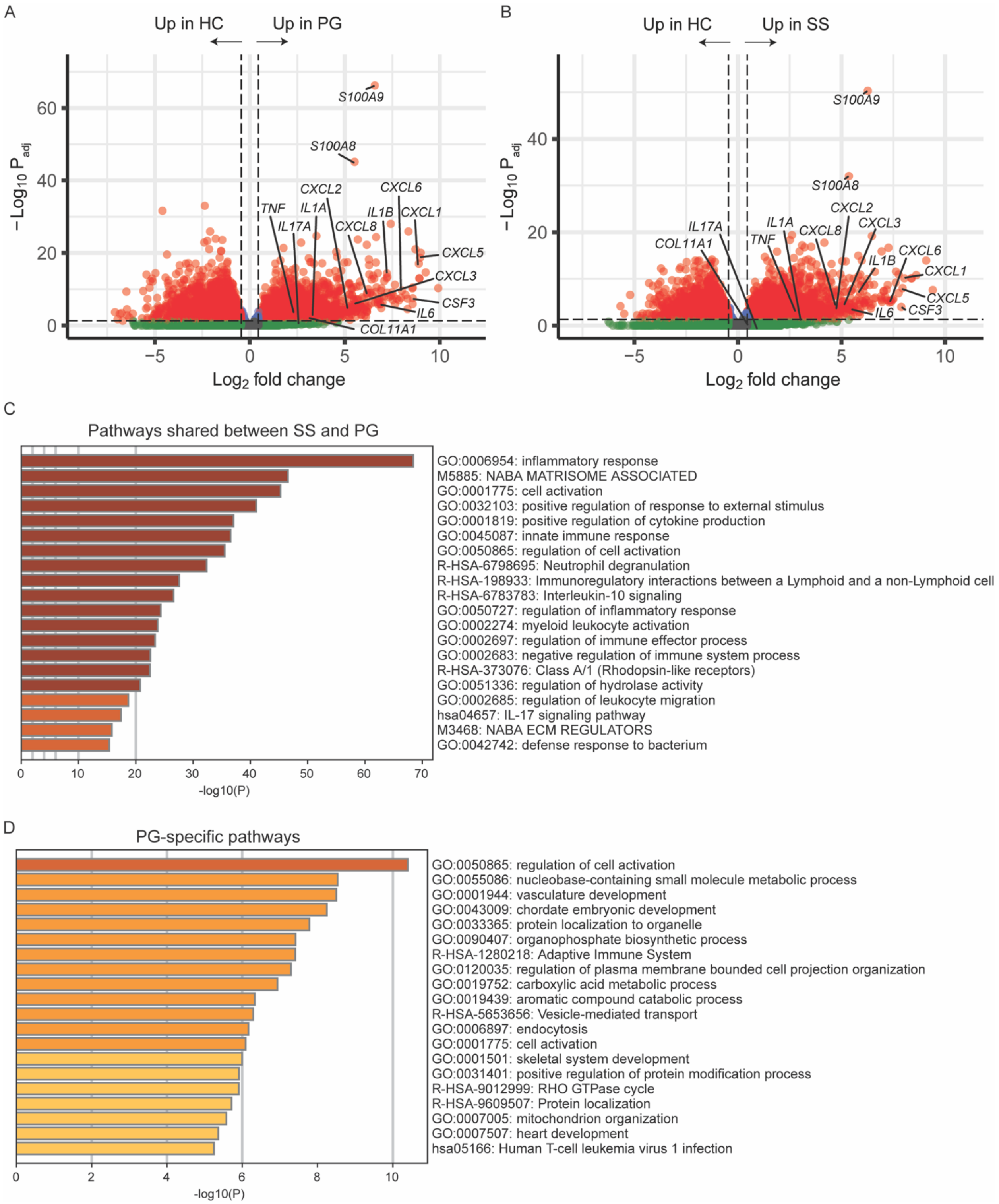


**Figure S4:** Sweet's syndrome and pyoderma gangrenosum bulk RNA-Seq. (A) Differentially expressed genes in PG compared to healthy control. (B) Differentially expressed genes in SS compared to HC. (A-B) The adjusted (adj) P value threshold is 0.05 and $\log_2$ fold-change (FC) threshold is 0.45. (C) Pathway analysis using significantly differentially expressed genes (up and down; $P_{adj} < 0.05$) shared between SS and PG. (D) Pathway analysis with differentially expressed genes unique to PG (up and down; $P_{adj} < 0.05$; $|FC| > 2$). Sweet's syndrome; PG, pyoderma gangrenosum; HC, healthy control.

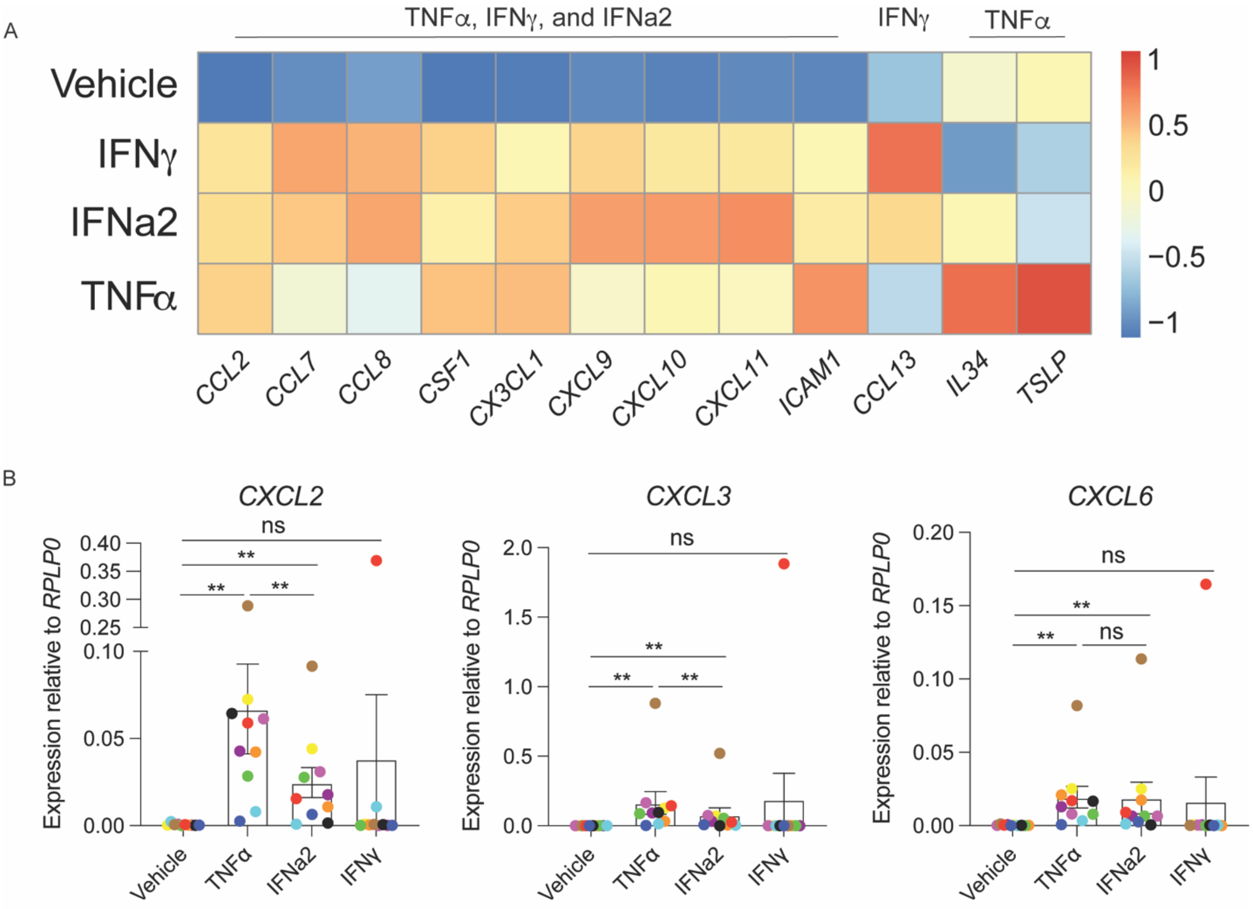


**Figure S5:** Bulk RNA-Seq of cultured primary human dermal fibroblasts. (A) Select inflammatory mediator genes highly upregulated by TNFα, IFNγ, and IFNa2; by IFNγ alone; or by TNFα alone. (B) qPCR gene expression of neutrophil chemokines. Each color represents a single donor's fibroblasts. ns (not significant) and **P < 0.01 using Wilcoxon matched pairs signed rank test.

A

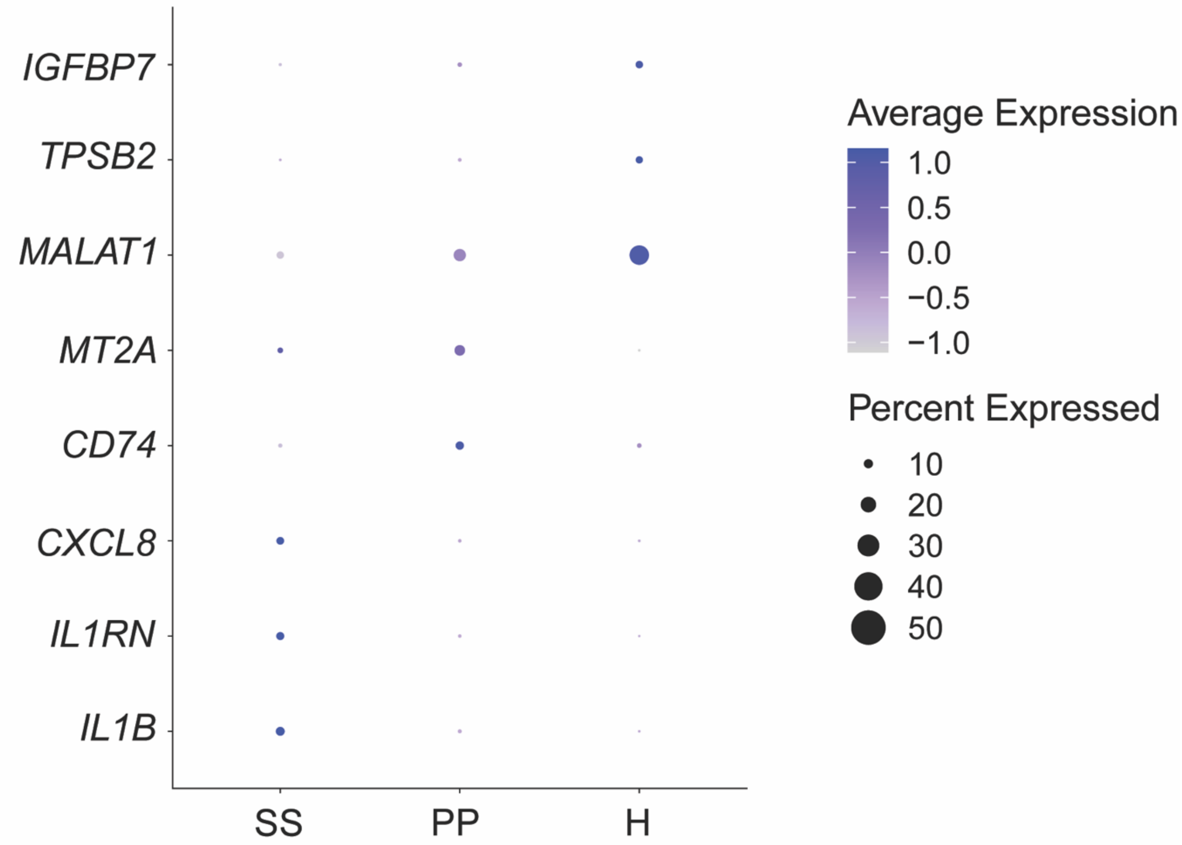


**Figure S6:** Neutrophils are hyperactivated in Sweet's syndrome. (A) Top 3 differentially expressed genes ($P_{adj} < 0.05$) between neutrophils in SS, PP, and HC using spatial transcriptomics data. SS, Sweet's syndrome; PP, pustular psoriasis; HC, healthy control.

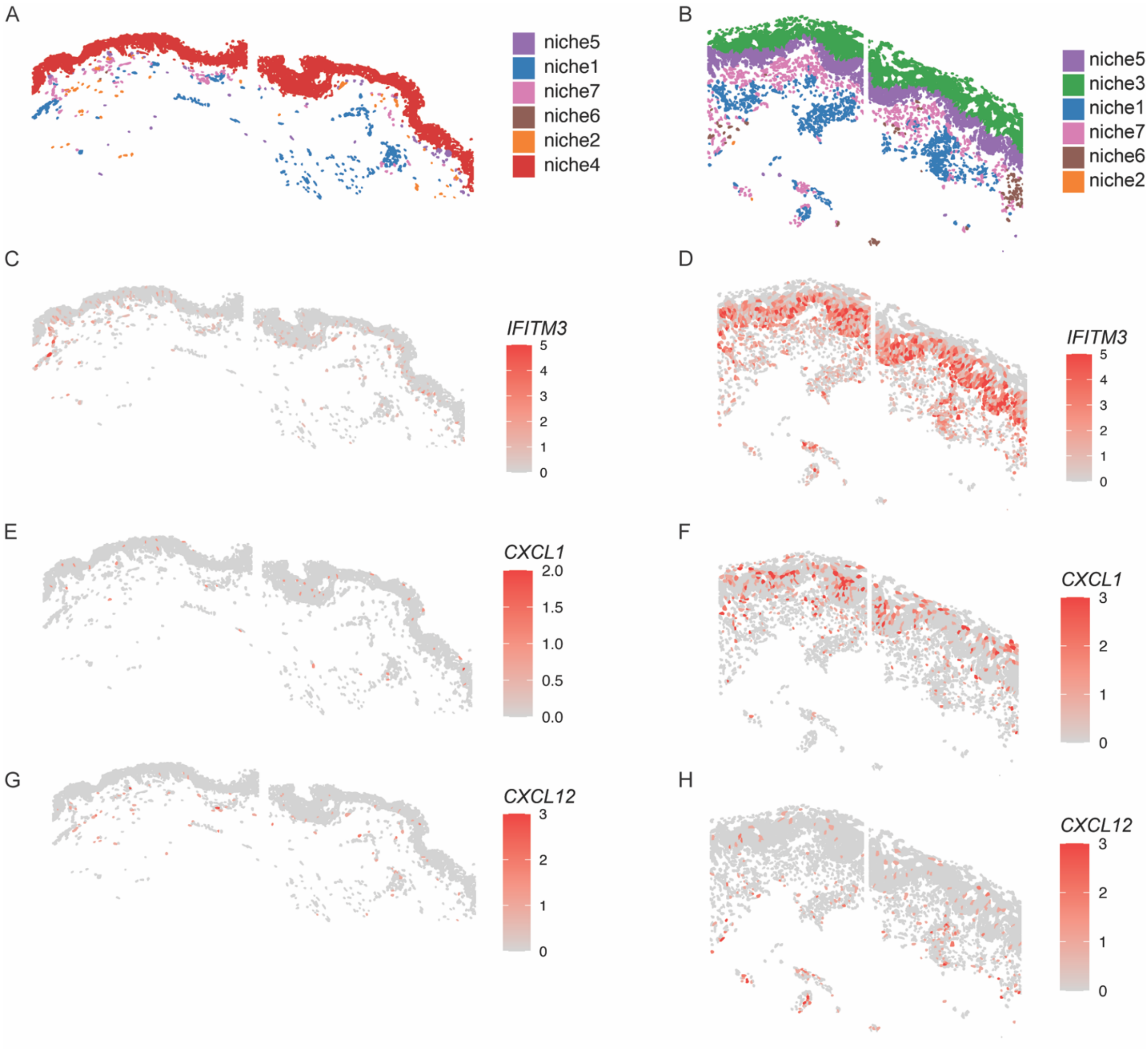


**Figure S7:** Spatial transcriptomic niche analysis of neutrophilic dermatoses. Niche analysis projected onto HC (A) and PP (B) tissue. HC expression of interferon-induced gene *IFITM3* (C) and neutrophil chemokines *CXCL12* (D) and *CXCL1* (E). SS, Sweet's syndrome; PG, pyoderma gangrenosum; PP, pustular psoriasis; HC, healthy control.
